# Canonical and non-canonical PRC1 differentially contribute to the regulation of neural stem cell fate

**DOI:** 10.1101/2024.08.07.606990

**Authors:** Janine Hoffmann, Theresa M. Schütze, Annika Kolodziejczyk, Annekathrin Kränkel, Susanne Reinhardt, Razvan P. Derihaci, Cahit Birdir, Pauline Wimberger, Haruhiko Koseki, Mareike Albert

## Abstract

Neocortex development is characterized by sequential phases of neural progenitor cell (NPC) expansion, neurogenesis and gliogenesis. Polycomb-mediated epigenetic mechanisms are known to play important roles in regulating the lineage potential of NPCs during development. The composition of Polycomb Repressive Complex 1 (PRC1) is highly diverse in mammals and was hypothesized to contribute to context-specific regulation of cell fate. Here, we have performed side-by-side comparison of the role of canonical PRC1.2/1.4 and non-canonical PRC1.3/1.5, all of which are expressed in the developing neocortex, in NSC proliferation and differentiation. We found that the deletion of *Pcgf2/4* in NSCs led to a strong reduction in proliferation and to altered lineage fate, both during the neurogenic and gliogenic phase, whereas *Pcgf3/5* played a minor role. Mechanistically, genes encoding stem cell and neurogenic factors were bound by PRC1 and differentially expressed upon *Pcgf2/4* deletion. Thus, rather than different PRC1 sub-complexes contributing to different phases of neural development, we found that canonical PRC1 played a more significant role in NSC regulation during proliferative, neurogenic and gliogenic phases compared to non-canonical PRC1.

## Introduction

During the development of the neocortex, stem and progenitor cells initially proliferate, before sequentially giving rise to neurons destined to different cortical layers and subsequently generate astrocytes and oligodendrocytes (Lodato & Arlotta, 2015; Qian *et al*, 2000). Precise spatial and temporal regulation of NPC proliferation and differentiation is key for the proper formation of the intricate structure of the neocortex. Transcription factors and epigenetic mechanisms play important roles in orchestrating dynamic changes in gene expression that underlie coordinated neural differentiation programs in the developing neocortex (Albert & Huttner, 2018; Desai & Pethe, 2020; Tsuboi *et al*, 2019).

Chromatin modifiers of the Trithorax and Polycomb groups maintain active and repressed gene activity states during embryonic development (Piunti & Shilatifard, 2016; Ringrose & Paro, 2004; Schuettengruber *et al*, 2017). Polycomb proteins assemble into two major complexes, PRC1 and PRC2, which catalyse mono-ubiquitination of histone 2A lysine 119 (H2AK119ub1) and tri-methylation of histone 3 lysine 27 (H3K27me3), respectively. These complexes are important determinants of the ability of NPCs to either proliferate or to give rise to neurons or glial cells (Hirabayashi *et al*, 2009; Tyssowski *et al*, 2014), and mutations in Polycomb components were reported to cause neurodevelopmental disorders (Bölicke & Albert, 2022; Mastrototaro *et al*, 2017; Pierce *et al*, 2018).

Specific deletion of the PRC2 histone methyltransferase *Ezh2* in the early developing neocortex causes an up-regulation of gene expression and a shift of apical radial glia fate from self-renewal to differentiation (Pereira *et al*, 2010), reducing the neuronal output and leading to a substantially smaller neocortex. Moreover, deletion of *Ring1b*, an integral component of PRC1, during the neurogenic phase results in altered neuronal subtype specification (Morimoto-Suzki *et al*, 2014). In this context, the E3 ubiquitin ligase activity of Ring1b was found to be necessary for the temporary repression of key neuronal genes in neurogenic NPCs (Tsuboi *et al*, 2018). These data indicate that Polycomb complexes control important aspects of corticogenesis.

Epigenome profiling in specific neural cell populations isolated from the developing mouse neocortex has revealed dynamic changes in H3K4me3 and H3K27me3 during neocortical lineage specification (Albert *et al*, 2017). An important question is how Polycomb target gene specificity is achieved in different neocortical cell types. One way to dynamically control Polycomb function and targeting is by altering the composition of Polycomb complexes, which in mammals is highly diverse, enabling the assembly of various sub-complexes with different functionalities (Kim & Kingston, 2020; Luis *et al*, 2012; Tsuboi *et al*., 2019). During neocortex development, chromatin remodelers of the chromodomain helicase DNA-binding (Chd) family, which interact with PRC2 complexes, show differential expression. Whereas Chd5 is expressed in neurons and controls neuronal differentiation (Egan *et al*, 2013), Chd4 is expressed in neural progenitor cells during early neurogenesis, where it functions in the inhibition of astroglial differentiation (Sparmann *et al*, 2013). Such a switch in subunit composition may contribute to the re-targeting of PRC2 during neocortex development.

PRC1 complexes are classified as canonical or non-canonical, depending on which of the complex members are included, with all complexes containing the central Ring1a/b E3 ubiquitin ligase core (Blackledge & Klose, 2021; Piunti & Shilatifard, 2016; Tsuboi *et al*., 2019). Canonical PRC1 complexes (PRC1.2/1.4) contain Pcgf2/4, one of three polyhomeotic (Phc) proteins and one of five chromobox (Cbx) proteins that recognize H3K27me3 mediated by PRC2. Non-canonical PRC1 is targeted to chromatin independently of H3K27me3 and is characterized by the inclusion of Pcgf1 (PRC1.1), Pcgf3/5 (PRC1.3/1.5) or Pcgf6 (PRC1.6) (Blackledge & Klose, 2021), even though non-canonical PRC1.2/1.4 lacking Cbx and Phc has also been described (Gao *et al*, 2012). Non-canonical PRC1 has high ubiquitin ligase activity, whereas canonical PRC1 was reported to promote higher order chromatin structures, but to display low ligase activity and low contribution to target gene repression (Blackledge & Klose, 2021; Fursova *et al*, 2019).

In embryonic stem cells, the interchange of Cbx family proteins in PRC1 (Kim & Kingston, 2020) has been reported to modulate the balance between self-renewal and lineage commitment (Morey *et al*, 2012; O’Loghlen *et al*, 2012; Santanach *et al*, 2017), and different Cbx paralogs are required for different cell lineages (Klauke *et al*, 2013; Luis *et al*, 2011). Likewise, Pcgf homologs were suggested to promote context- and stage-specific functions during differentiation and development (Kloet *et al*, 2016; Morey *et al*, 2015).

While the canonical PRC1 components Pcgf2 and Pcgf4 (Akasaka *et al*, 2001; Fasano *et al*, 2007; He *et al*, 2009; Leung *et al*, 2004; Molofsky *et al*, 2005; Molofsky *et al*, 2003; Zencak *et al*, 2005) as well as the non-canonical PRC1 components Pcgf3 and Pcgf5 (Gao *et al*, 2014; Meng *et al*, 2020; Yao *et al*, 2018) have been implicated in neural differentiation and brain development, here we set out to perform a systemic comparative analysis of the role of different PRC1 subcomplexes in NSC proliferation and differentiation. Specifically, we deleted canonical (*Pcgf2/4*) and non-canonical PRC1 (*Pcgf3/5*) in proliferating, neurogenic and gliogenic NSCs to elucidate the function of different PRC1 sub-complexes in key phases of cortical development.

## Results and discussion

### Pcgf homologs are differentially expressed in the mouse and human developing neocortex

To analyse the expression of canonical and non-canonical PRC1 components (Figure 1A) in the human developing neocortex (Figure 1B), we first mined RNA-seq data of microdissected human foetal cortex (Fietz *et al*, 2012). The core components of PRC1, *RING1A* and *RING1B*, showed a fairly uniform distribution across the germinal zones (VZ, ISVZ, OSVZ), which are enriched in NPCs, and the cortical plate, where neurons reside (Figure 1C). This was confirmed by immunohistochemistry of human foetal tissue, in which RING1B and H2AK119ub1 showed a comparable expression across all cortical layers with a minor increase in the ventricular zone (VZ) and cortical plate (CP) (Figure 1D, E). In contrast, *PCGF2*, *PCGF3* and *PCGF4* were expressed at higher levels in the CP compared to germinal zones, whereas *PCGF5* was specifically enriched in the VZ (Figure 1C–E). These data indicate that PCGF homologs are differentially expressed across neural cell types of the human developing neocortex.

**Figure 1.**
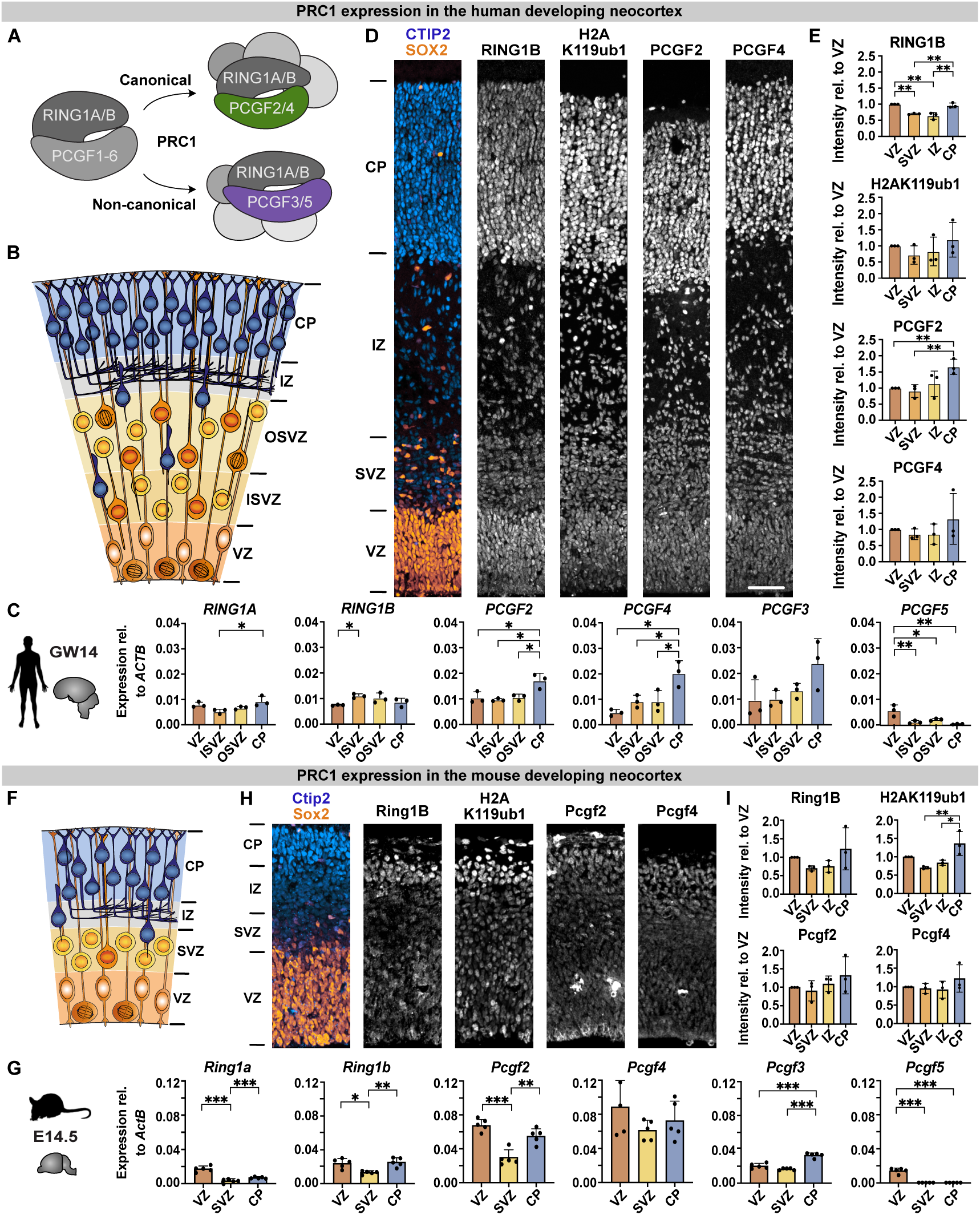
PRC1 components are differentially expressed in the human and mouse developing neocortex. (**A**) Schematic illustration of canonical and non-canonical PRC1 complexes containing the RING1A/B core and either PCGF2/4 or PCGF3/5 subunits, respectively. (**B**) Schematic illustration of the human developing neocortex, divided into the ventricular zone (VZ), inner subventricular zone (ISVZ), outer subventricular zone (OSVZ), intermediate zone (IZ) and cortical plate (CP). (**C**) PRC1 mRNA expression levels in the human developing neocortex at gestation week (GW) 14 analysed by RNA-seq (data from Fietz *et al*. (2012)), relative to the house keeping gene *ACTB*. (**D**) Immunofluorescence for the radial glia marker SOX2, neuronal marker CTIP2, and PRC1-related RING1B, H2AK119ub1, PCGF2 and PCGF4 of human foetal tissue (GW12/13). (**E**) Quantifications of mean intensity per cell in the indicated zones of human foetal tissue, relative to the intensity in the VZ. (**F**) Schematic illustration of the mouse developing neocortex. (**G**) PRC1 mRNA expression levels in the mouse developing neocortex at E14.5 analysed by RNA-seq (data from Fietz *et al*. (2012)), relative to the house keeping gene *Actb*. (**H**) Immunofluorescence of mouse embryonic tissue (E14.5). (**I**) Quantifications of mean intensity per cell in the indicated zones of mouse embryonic tissue, relative to the intensity in the VZ. Data information: Scale bars, 100 µm. Bar graphs represent mean values. Error bars represent SD; C, D, of 3 tissue samples from different individuals; G, I, of 3–5 embryos from at least two different litters. *** *p* < 0.001, ** *p* < 0.01, * *p* < 0.05; Tukey’s multiple comparison test.

Next, we analysed expression of PRC1 components in the mouse developing neocortex (Figure 1F). Ring1a/b and H2AK119ub1 were uniformly distributed across all zones, with a slight enrichment in the VZ and CP (Figure 1G–I). In analogy to the human developing neocortex, Pcgf2, *Pcgf3* and Pcgf4 showed some enrichment in the CP, whereas *Pcgf5* was specifically expressed in the VZ (Figure 1G–I), even though the differences were less pronounced in the mouse compared to the human developing neocortex. The results are in line with previous studies noting the high abundance of Pcgf2 and Pcgf4 in NPCs (Tagawa *et al*, 1990; Zencak *et al*., 2005), whereas neuronal expression of Pcgf4 was not seen previously (Zencak *et al*., 2005).

Taken together, the canonical PRC1 components Pcgf2/4 were enriched in neurons compared to NPCs, which is interesting giving their previous implication in the regulation of NSC self-renewal and proliferation (Fasano *et al*., 2007; He *et al*., 2009; Molofsky *et al*., 2005; Molofsky *et al*., 2003; Zencak *et al*., 2005). The non-canonical PRC1 components Pcgf3 and Pcgf5 showed differential enrichment in neurons versus NPCs, respectively.

### Canonical and non-canonical PRC1 contribute to different degrees to the regulation of NSC proliferation

To systematically compare the role of canonical and non-canonical PRC1 in NSC proliferation, we isolated NPCs from the developing dorsolateral neocortex of E12.5 mouse embryos from either *Pcgf2*^F/F^; *Pcgf4*^F/F^; *Nes*::CreERT2/+ or *Pcgf3*^F/F^; *Pcgf5*^F/F^; *Nes*::CreERT2/+ mouse lines (Almeida *et al*, 2017; Fursova *et al*., 2019; Imayoshi *et al*, 2006) and induced the deletion of *Pcgf* genes *in vitro* by the addition of 4-hydroxytamoxifen (OHT) (Figure 2A). Deletion of *Pcgf* genes was highly efficient (Figure 2B, C) and resulted in a complete loss of Pcgf3, Pcgf4 and Pcgf5 proteins after 3 days *in vitro* (DIV), and a reduction in Pcgf2 levels (Figure 2D, E).

**Figure 2.**
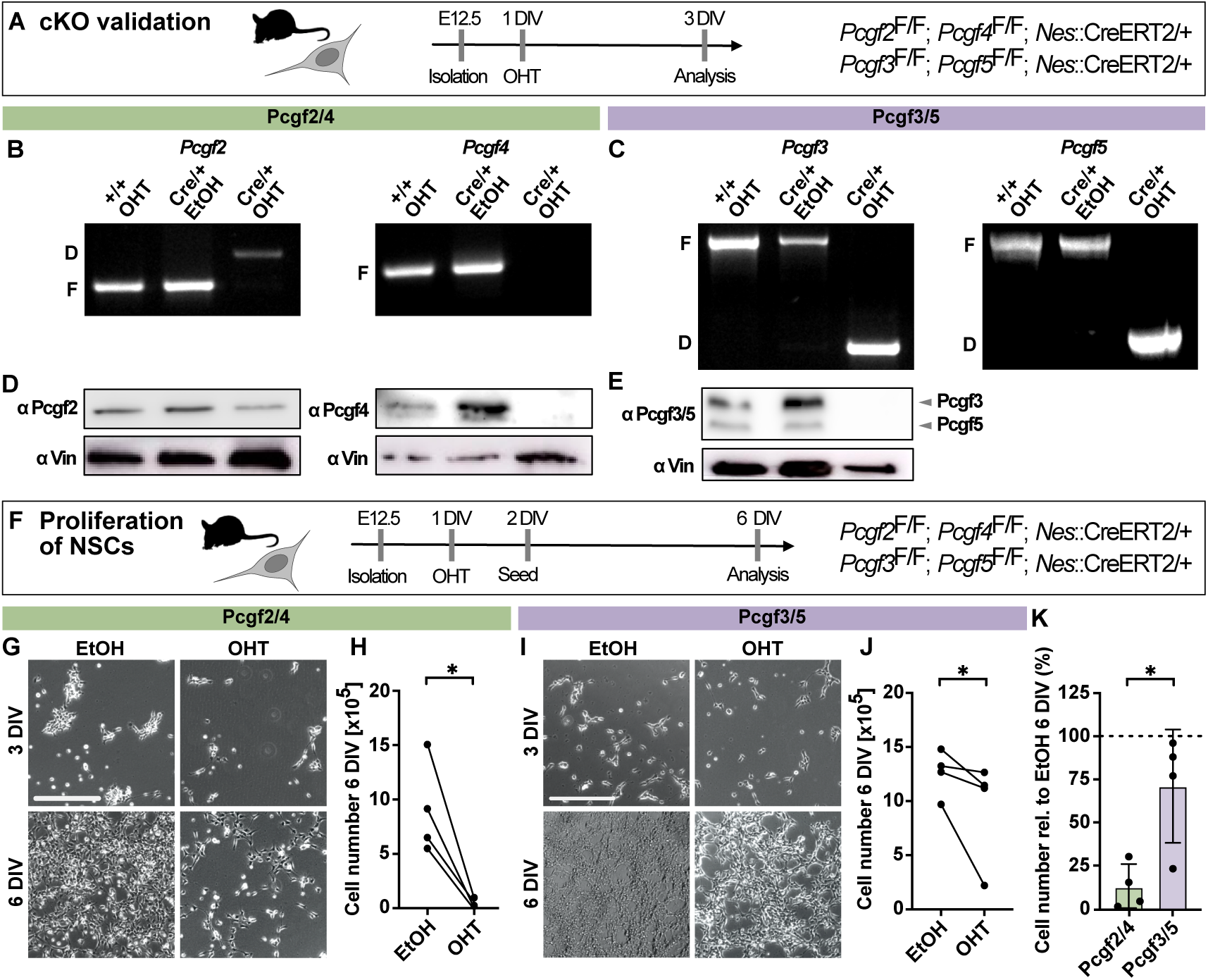
Deletion of *Pcgf2/4* and *Pcgf3/5* reduces NSC proliferation to different degrees. (**A**) Schematic of experimental workflow. NSCs were isolated from E12.5 embryos from either *Pcgf2*^F/F^; *Pcgf4*^F/F^; *Nes*::CreERT2/+ or *Pcgf3*^F/F^; *Pcgf5*^F/F^; *Nes*::CreERT2/+ mouse lines. After one day *in vitro* (DIV), the deletion of Pcgf components was induced by the administration of 4-hydroxytamoxifen (OHT) and analysis was performed at 3 DIV. (**B, C**) Genotyping of *Pcgf2/4* (B) and *Pcgf3/5* (C) floxed (‘F’) and deletion (‘D’) alleles by PCR analysis following treatment of NSC cultures from control (+/+) or experimental (Cre/+) mice with ethanol (‘EtOH’; control) or OHT. (**D, E**) Immunoblots of protein lysates from the same NSC cultures shown in (B), using anti-Pcgf2, anti-Pcgf4, anti-Pcgf3/5 and anti-Vinculin antibodies. (**F**) Schematic of experimental workflow. Deletion of *Pcgf* genes was induced at 1 DIV, 50,000 cells were seeded at 2 DIV and cells were counted at 6 DIV. (**G**) Brightfield images of *Pcgf2*^F/F^; *Pcgf4*^F/F^; *Nes*::CreERT2/+ NSC cultures treated with EtOH or OHT at 3 DIV and 6 DIV. (**H**) Quantification of cell numbers at 6 DIV. (**I**) Brightfield images of *Pcgf3*^F/F^; *Pcgf5*^F/F^; *Nes*::CreERT2/+ NSC cultures treated with EtOH or OHT at 3 DIV and 6 DIV. (**J**) Quantification of cell numbers at 6 DIV. (**K**) Data from (H, J) plotted for comparison of cell numbers following deletion of either *Pcgf2/4* or *Pcgf3/5*, shown as percentage relative to the EtOH control. Data information: Scale bars, 300 µm. Bar graphs represent mean values. Error bars represent SD. H, J, dots connected by lines represent 4 embryos from at least two independent litters treated with either EtOH or OHT. * *p* < 0.05; Tukey’s multiple comparison test.

Following the validation of conditional knockout (cKO) in NSCs, we then asked how deletion of canonical (*Pcgf2/4*) and non-canonical (*Pcgf3/5*) PRC1 affects NSC proliferation. Whereas deletion of *Pcgf2/4* led to an almost complete inability of NSCs to proliferate (Figure 2G, H), the deletion of *Pcgf3/5* had a more modest effect (Figure 2I, J). Overall, the cell numbers were reduced to less than 10% of control after 6 DIV for *Pcgf2/4* cKO compared to roughly 70% of control for *Pcgf3/5* cKO (Figure 2K), highlighting the differential contribution of canonical and non-canonical PRC1 to the regulation of NSC proliferation.

These results are in agreement with previous reports on the role of *Pcgf4* in regulating NSC self-renewal and proliferation (Fasano *et al*., 2007; He *et al*., 2009; Molofsky *et al*., 2005; Molofsky *et al*., 2003; Zencak *et al*., 2005). Pcgf2 was reported to function antagonistically to Pcgf4 by promoting cell senescence through down-regulation of Pcgf4 (Guo *et al*, 2007). Yet, double knockout of *Pcgf2/4* revealed the synergistic effect of both genes, resulting in strongly exacerbated phenotypes compared to single mutant mice (Akasaka *et al*., 2001). In line with this, deletion of *Pcgf2/4* in NSCs resulted in an almost complete loss of the ability of NSCs to proliferate. Moreover, *Pcgf3* has previously been implicated in the regulation of tumour cell proliferation (Hu *et al*, 2021), whereas *Pcgf5* and *Pcgf3*/*5* were dispensable for embryonic stem cell self-renewal (Yao *et al*., 2018; Zhao *et al*, 2017). Here, we showed that *Pcgf3/5* regulate NSC proliferation.

In summary, side-by-side comparison of double knockout of different *Pcgf* homologs revealed a stronger contribution of canonical PRC1.2/1.4 compared to non-canonical PRC1.3/1.5 to the regulation of NSC proliferation.

### Canonical, but not non-canonical, PRC1 regulates the differentiation potential of NSCs

Next, we aimed to dissect the contribution of canonical and non-canonical PRC1 to NSC differentiation. For this, we made use of the previously described potential of NSCs to maintain their developmental progression *in vitro*, initially resulting in the production of neurons (neurogenic phase), followed by the generation of astrocytes (gliogenic phase) (Hirabayashi *et al*., 2009). Deletion of *Pcgf2/4* in freshly isolated NSCs, that were induced to differentiate by the withdrawal of growth factors and the addition of serum to the medium, resulted in the generation of more neurons at the expense of oligodendrocytes, leaving the proportion of astrocytes unchanged (Figure 3A–C). In contrast, deletion of *Pcgf3/5* did not result in any significant changes in the proportions of differentiated cell types. This suggests that canonical and non-canonical PRC1 complexes differentially contribute to the regulation of NSC fate during the neurogenic phase.

**Figure 3.**
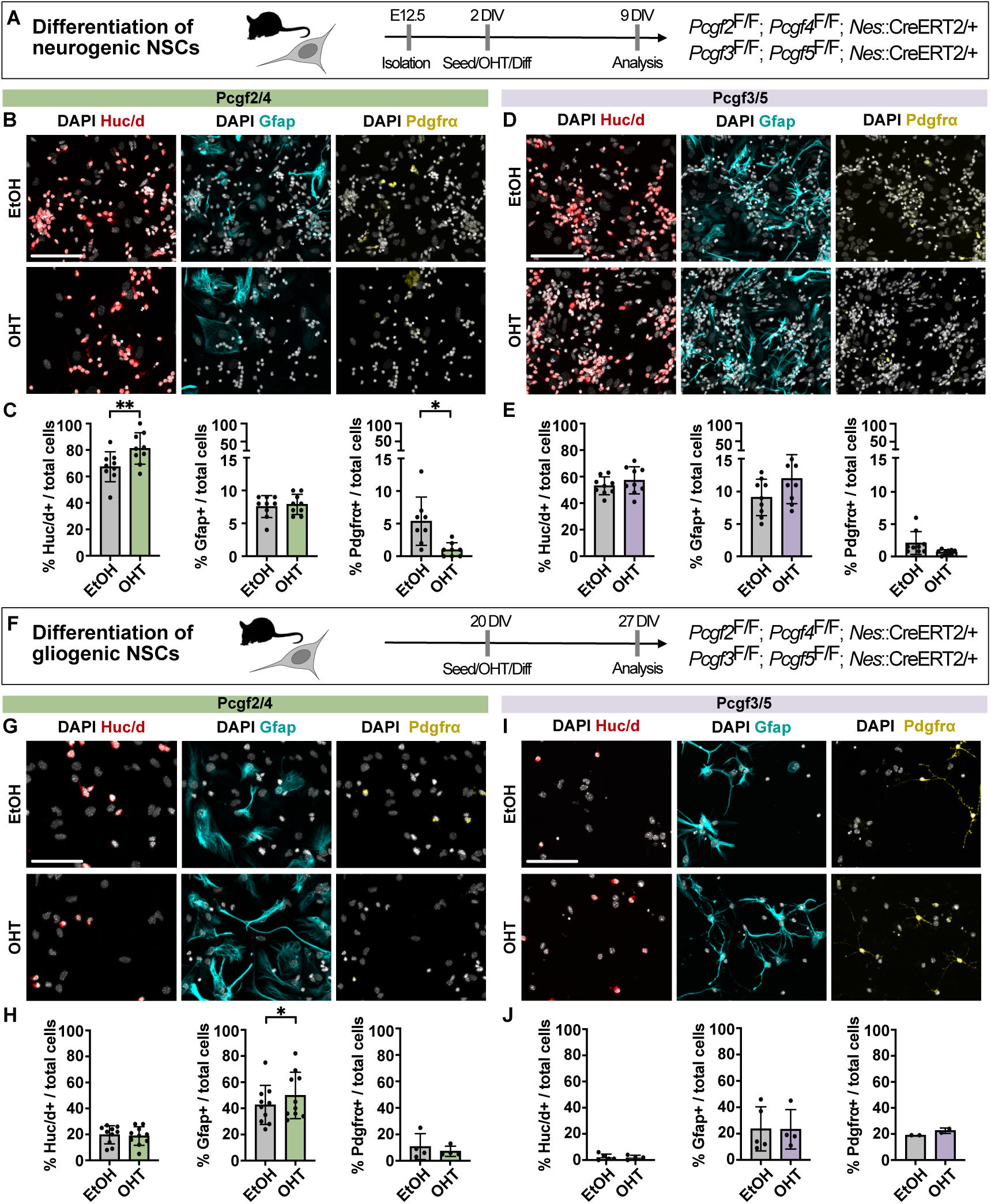
Deletion of *Pcgf2/4* but not *Pcgf3/5* results in altered lineage potential of neurogenic and gliogenic NSCs. (**A**) Schematic of experimental workflow. Deletion of *Pcgf* genes was induced at 2DIV concomitant with seeding of NSCs with neurogenic potential and induction of differentiation. Differentiated cells were analysed at 9 DIV. (**B**) DAPI staining and immunofluorescence for the pan-neuronal marker Huc/d, the astrocyte marker Gfap and the oligodendrocyte precursor marker Pdgfrα after 7 days of differentiation of *Pcgf2*^F/F^; *Pcgf4*^F/F^; *Nes*::CreERT2/+ NSCs. (**C**) Quantification of the percentage of marker-positive cells of total DAPI-positive cells following deletion of *Pcgf2/4*. (**D**) DAPI staining and immunofluorescence after 7 days of differentiation of *Pcgf3*^F/F^; *Pcgf5*^F/F^; *Nes*::CreERT2/+ NSCs. (**E**) Quantification of the percentage of marker-positive cells of total DAPI-positive cells following deletion of *Pcgf3/5*. (**F**) Schematic of experimental workflow. Deletion of *Pcgf* genes was induced at 20 DIV in NSCs with gliogenic potential, concomitant with seeding and induction of differentiation. (**G**) DAPI staining and immunofluorescence for lineage markers. (**H**) Quantification of the percentage of marker-positive cells. (**I**) DAPI staining and immunofluorescence for lineage markers. (**J**) Quantification of the percentage of marker-positive cells. Data information: Scale bars, 100 µm. Bar graphs represent mean values. Error bars represent SD; C, E, H, J, of 4–9 embryos from at least two different litters. ** *p* < 0.01, * *p* < 0.05; Tukey’s multiple comparison test.

To compare the contribution of different PRC1 subcomplexes during the gliogenic phase, we kept the NSC lines in culture for 20 days and then repeated the differentiation experiment (Figure 3F). While cKO of *Pcgf2/4* resulted in the generation of more astrocytes compared to control, cKO of *Pcgf3/5* again did not affect the differentiation potential of NSCs. This side-by-side comparison highlights the role of canonical PRC1.2/1.4, but not non-canonical PRC1.3/1.5, in determining the linage potential of NSCs during differentiation.

Taken together, deletion of *Pcgf2/4* resulted in an increased proportion of neurons during the neurogenic phase and of astrocytes during the gliogenic phase. This is in contrast to the deletion of *Ring1b*, central to all PRC1 complexes, which was shown to not affect neuron numbers during the neurogenic phase, but resulted in more neurons during the gliogenic phase (Hirabayashi *et al*., 2009). These results underscore the differential contributions of PRC1 subcomplexes with different subunit composition during neurogenic and gliogenic phases of neural differentiation.

### Canonical PRC1 regulates the expression of stem cell and neurogenic factors

Lastly, to explore the mechanism by which canonical PRC1 regulates NSC fate during differentiation, we performed gene expression analysis by RNA-seq, 4 days after induction of differentiation, following deletion of *Pcgf2/4* in neurogenic NSCs (Figure 4A, B). In line with the repressive function of PRC1, we identified 120 genes that were more than 2-fold up-regulated upon deletion of *Pcgf2/4* compared to control, whereas only 24 genes were down-regulated (Figure 4C). The differentially expressed genes (DEG) were characterized by gene ontology (GO) terms related to the molecular functions ‘DNA-binding’ and ‘E-box-binding’ (Figure 4D), which is in accordance with hallmarks of Polycomb-mediated regulation through binding of genes encoding key developmental transcription factors (Bernstein *et al*, 2006; Schuettengruber *et al*., 2017). Moreover, GO terms related to biological processes included ‘regionalization’ and ‘pattern specification’ (Figure 4E), extending what has been described for other Polycomb proteins during brain development (Albert *et al*., 2017; Eto *et al*, 2020; Hirabayashi *et al*., 2009). Of the 120 up-regulated genes, the majority (89 genes) were directly bound by Pcgf2 and/or Ring1b in NPCs (Figure 4F) (Kloet *et al*., 2016). Among the direct PRC1 targets that were up-regulated, we found many transcriptions factors (Figure 4G), including *Hox* genes (Figure 4H), which represent known targets of PRC1 (Akasaka *et al*., 2001; Kloet *et al*., 2016).

**Figure 4.**
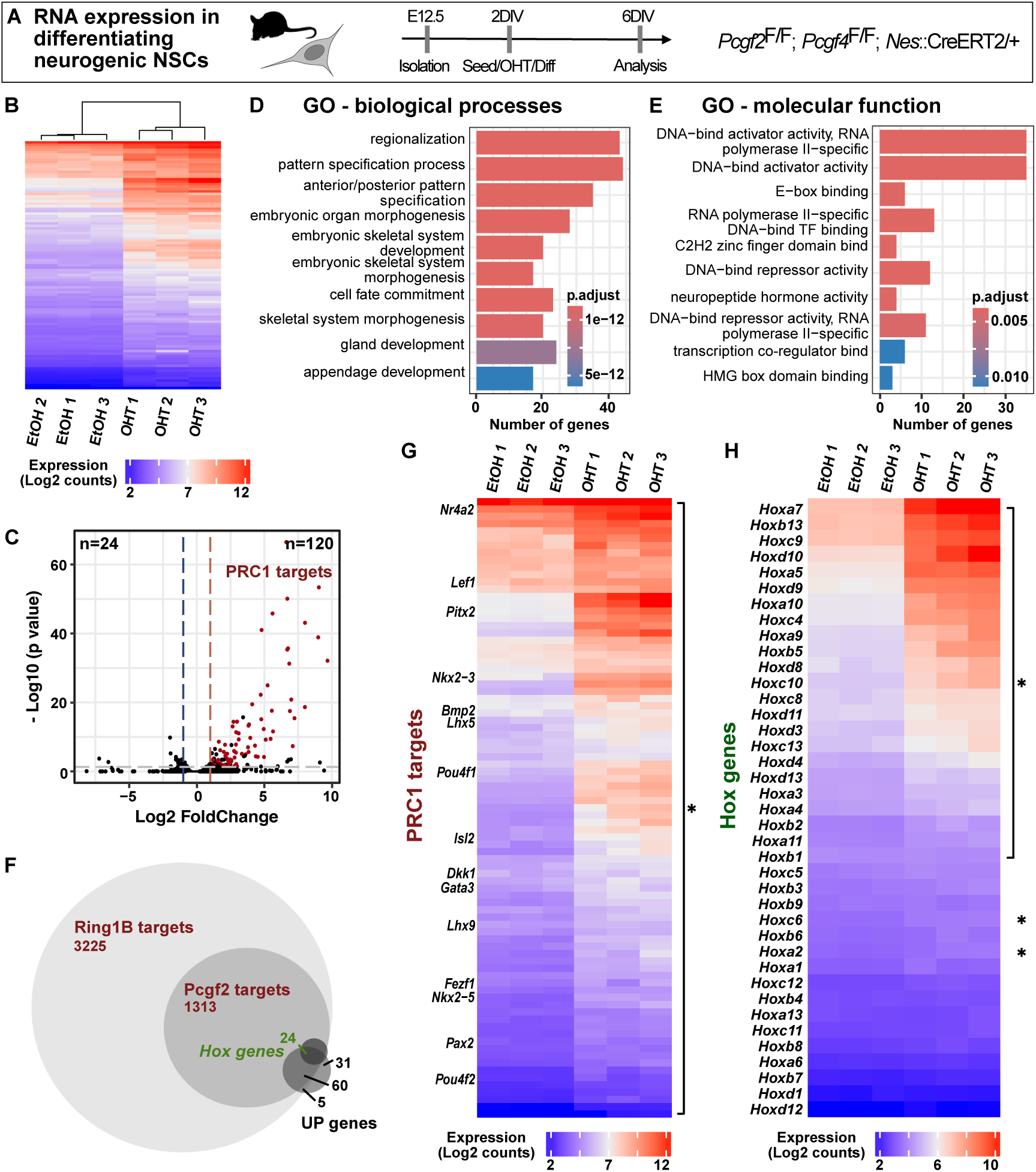
Deletion of *Pcgf2/4* results in derepression of PRC1 target genes linked to fate specification. (**A**) Schematic of experimental workflow. Deletion of *Pcgf2/4* genes was induced at 2 DIV concomitant with seeding of NSCs with neurogenic potential and induction of differentiation. Gene expression was analysed by RNA-seq after 4 days of differentiation at 6 DIV. (**B**) Hierarchical clustering analysis with the heatmap of the 100 most differentially expressed genes between control (EtOH) and *Pcgf2/4* deletion (OHT) samples, showing the clustering of replicates. (**C**) Volcano plot of log10 (*p* value) against log2 fold change representing the differences in gene expression between control samples (EtOH) and *Pcgf2/4* cKO (OHT). Grey line represents cutoff of *p* < 0.05, blue line of log2 fold change < −1 and red line of log2 fold change > 1. Numbers in the upper corners indicate number of significantly up- and down-regulated genes, respectively. Genes bound by Ring1b and/or Pcgf2 in NPCs (Kloet *et al*., 2016) are highlighted (‘PRC1 targets’). (**D, E**) Gene ontology (GO) term enrichment analysis of upregulated genes (*p* value < 0.05) was performed for biological processes (**D**) and molecular function (**E**). (**F**) Venn diagram representing the overlap of genes bound by Ring1b and Pcgf2 in NPCs (Kloet *et al*., 2016) with up-regulated genes, which include several *Hox* genes. (**G**) Heat map of the genes up-regulated following *Pcgf2/4* deletion (OHT) and bound by Ring1b and Pcgf2 in NPCs (Kloet *et al*., 2016), with a log2 fold change >1. (**H**) Heat map of up-regulated *Hox* genes, with a log2 fold change >1. Data information: Replicates represent NSCs from 3 embryos from two different litters.

Next, we aimed to specifically explore the expression of factors that may underlie the shifts in NSC lineage potential that we observed in *Pcgf2/4* cKO NSCs. We found that several stem cell factors, including *Id2*, *Pax6* and *Hes5*, showed reduced expression after *Pcgf2/4* deletion, whereas neurogenic factors (*Lhx5*, *Lhx9*, *Nr4a2*) and neuronal maturation genes (*En1*, *En2*, *Pitx3*) were increased compared to control (Figure 5A–D). These genes were bound by Pcgf2 and/or Ring1b in NPCs (Kloet *et al*., 2016), suggesting that they may represent direct targets of canonical PRC1 and may contribute to the enhanced neuronal differentiation of neurogenic NSCs after deletion of *Pcgf2/4*.

**Figure 5.**
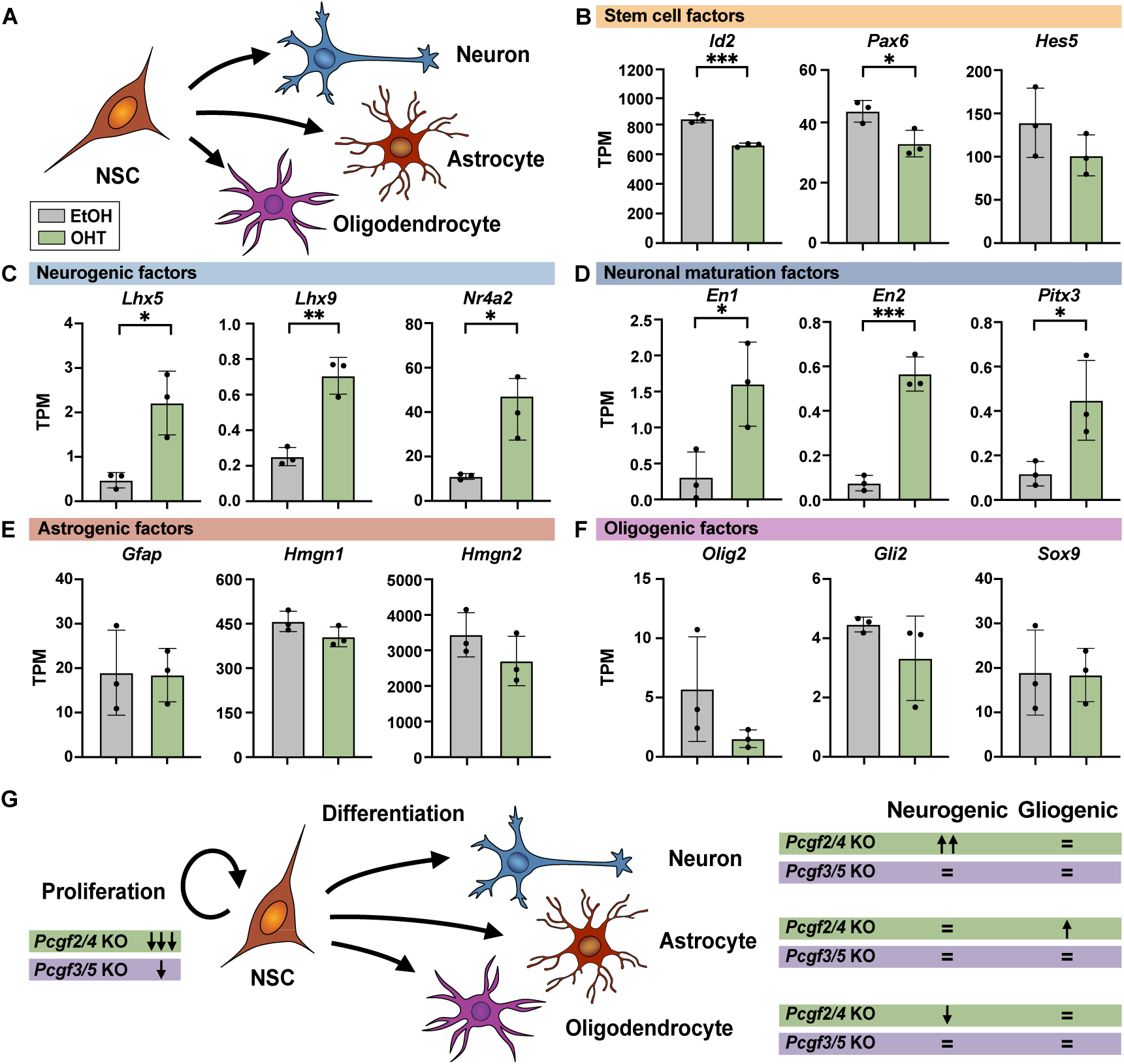
Deletion of *Pcgf2/4* affects expression of stem cell and neurogenic genes. (**A**) Schematic of NSC differentiation paradigm. (**B–F**) mRNA expression in transcripts per million (TPM) analysed by RNA-seq for genes encoding (A) stem cell, (B) neurogenic, (C) neuronal maturation, (E) astrogenic and (F) oligogenic factors. (**G**) Summary of the functional role of *Pcgf2/4* and *Pcgf3/5* in the regulation of NSC proliferation and differentiation during the neurogenic and gliogenic phase. Data information: Bar graphs represent mean values. Error bars represent SD, NSCs from 3 embryos from two independent litters. *** *p* < 0.001, ** *p* < 0.01, * *p* < 0.05; unpaired Student’s *t*-test.

In contrast, regulators of astrocyte fate, such as *Gfap*, *Hmgn1* and *Hmgn2*, did not show altered expression (Figure 5E) and were not bound by Polycomb (Albert *et al*., 2017; Kloet *et al*., 2016), which is in agreement with previous reports suggesting that astrocyte-specific genes are regulated by DNA methylation in NPCs (Fan *et al*, 2005; Hatada *et al*, 2008). Several genes encoding oligogenic factors (*Olig2*, *Gli2*, *Sox9*) were directly bound by Pcgf2 and/or Ring1b in NPCs (Kloet *et al*., 2016) and showed a trend for reduced expression, even though not significantly (Figure 5F).

In summary, our side-by-side comparison of *Pcgf2/4* and *Pcgf3/5* deletion in NSCs revealed a differential contribution of canonical and non-canonical PRC1, respectively, to the regulation of NSC proliferation and lineage potential (Figure 5G). Despite the observation that Pcgf2/4 expression is highest in the neuronal population, we observed that canonical PRC1.2/1.4 contributes to the regulation of NSCs at proliferative, neurogenic and gliogenic phases. Even though *Pcgf5* is preferentially expressed in the ventricular zone *in vivo*, deletion of *Pcgf3/5* only had a minor effect on NSC proliferation *in vitro* and did not impact NSC differentiation, neither during the neurogenic nor the gliogenic phase. It remains possible that additional non-canonical complexes (PRC1.1/1.6) may functionally contribute to the regulation of NSC fate, even though at least *Pcgf1* is expressed at low levels in the mouse developing neocortex (Fietz *et al*., 2012).

Mechanistically, PRC1 was reported to bind to several stem cell and neurogenic genes, which we found to be differentially expressed upon *Pcgf2/4* deletion, suggesting that these genes might be directly regulated by canonical PRC1. Our data suggest that rather than different PRC1 sub-complexes contributing to different phases of neural development, it is canonical PRC1.2/1.4 that regulates NSC proliferation and differentiation, whereas PRC1.3/1.5 plays a minor role in these processes. Overall, this suggests that switches in subunit composition may be more characteristic to the exit from pluripotency and differentiation towards different tissues and organs (Klauke *et al*., 2013; Luis *et al*., 2011; Morey *et al*., 2012; Morey *et al*., 2015; O’Loghlen *et al*., 2012; Santanach *et al*., 2017), but may be less relevant within a given lineage, such as neural development.

## Materials and methods

### Reagents and tools table

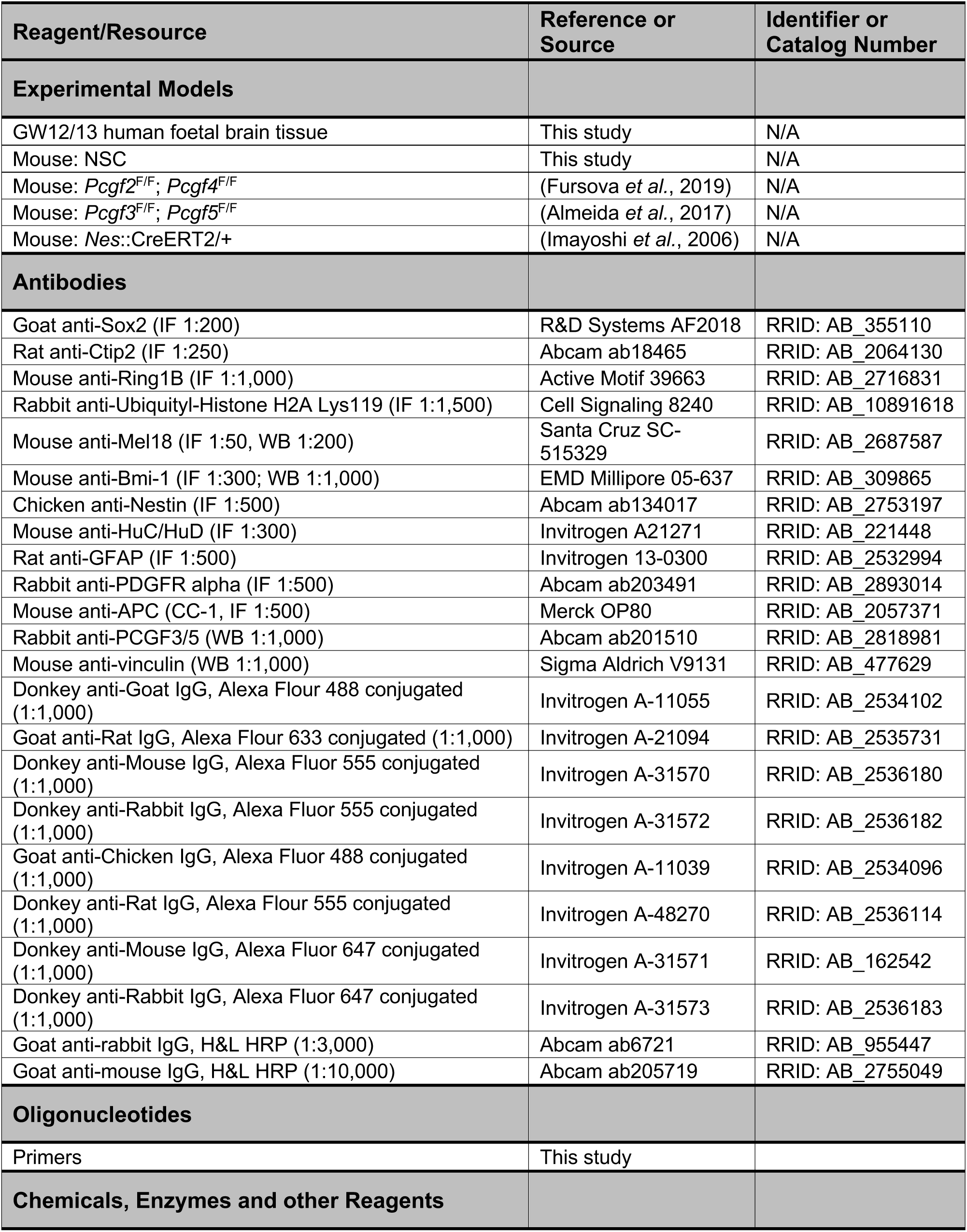
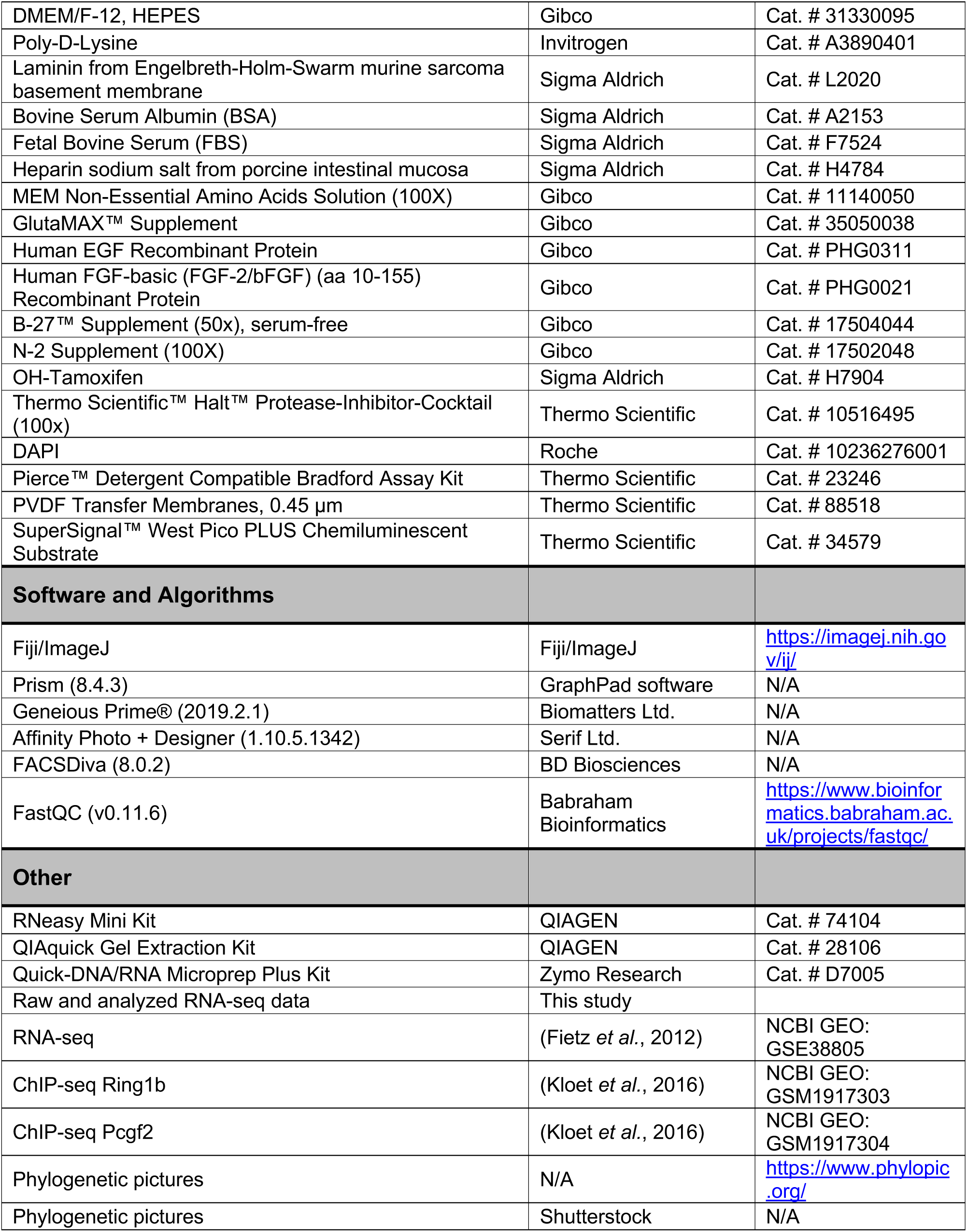

### Methods

#### Mice

All experimental procedures were conducted in agreement with the German Animal Welfare Legislation after approval by the Landesdirektion Sachsen (licenses DD24.1-5131/476/8; 25-5131/521/16). Animals were kept on a 12-hour/12-hour light/dark cycle with food and water *ad libitum*. Mice used for PRC1 expression analysis were wildtype mice from the inbred C57BL/6J strain. Embryonic day (E) 0.5 was set as noon on the day on which the vaginal plug was observed. All experiments were performed in the dorsolateral telencephalon of mouse embryos, at a medial position along the rostro-caudal axis. The developmental time point E14.5 of experimental procedures corresponds to a mid-neurogenic stage, when the production of upper-layer neurons has started. The sex of embryos was not determined, as it is not likely to be of relevance for the results obtained in the present study.

For the inducible deletion of *Pcgf* genes in NSCs, *Pcgf2*^F/F^; *Pcgf4*^F/F^ (Fursova *et al*., 2019) and *Pcgf3*^F/F^; *Pcgf5*^F/F^ (Almeida *et al*., 2017) mice were crossed with *Nes*::CreERT2/+ mice (Imayoshi *et al*., 2006). NSCs were isolated from either *Pcgf2*^F/F^; *Pcgf4*^F/F^; *Nes*::CreERT2/+ or *Pcgf3*^F/F^; *Pcgf5*^F/F^; *Nes*::CreERT2/+ strains. Only male embryos were included for *Pcgf3*^F/F^; *Pcgf5*^F/F^; *Nes*::CreERT2/+ to exclude effects attributed to changes in X chromosome inactivation (Almeida *et al*., 2017).

#### Human foetal brain tissue

Human foetal brain tissue was obtained from the Department of Gynaecology and Obstetrics, University Clinic Carl Gustav Carus of the Technische Universität Dresden, following elective pregnancy termination and informed written maternal consents, and with approval of the local University Hospital Ethical Review Committee (IRB00001473; IORG0001076; ethical approval number EK 355092018), in accordance with the Declaration of Helsinki. The age of foetuses ranged from gestation week (GW) 12 to 13 as assessed by ultrasound measurements of crown-rump length and other standard criteria of developmental stage determination. The developmental time point corresponds to an early/mid-neurogenic stage, when the OSVZ expands and the production of upper-layer neurons starts. Due to protection of data privacy, the sex of the human foetuses, from which the neocortex tissue was obtained, cannot be reported. The sex of the human foetuses is not likely to be of relevance for the results obtained in the present study. The foetal neocortex tissue samples used in this study reported no health disorders. Foetal human brain tissue was dissected in Tyrode’s solution and fixed immediately (within 1 hour).

#### Mouse NSC culture and differentiation

Mouse NSCs were isolated from E12.5 embryos as previously described (Cubillos *et al*, 2024; Schmitz *et al*, 2011; Schütze *et al*, 2022). Briefly, the dorsolateral cortex was isolated from E12.5 mice and treated with trypsin-EDTA (Gibco, #25300054) at 37°C for 20 min. Subsequently, soybean trypsin inhibitor (0.25mg/ml in PBS, Invitrogen, #17075-029) was added, cells were mechanically dissociated by pipetting and pelleted at 300x g for 5 min. NSCs were then plated on poly-D-lysine (at least 2 hours at 37°C; Gibco, #A3890401) and laminin (at least 4 hours at 37 °C; Sigma Aldrich, #L2020) coated plates at a density of 40,000 cells/mm^2^ and cultured under standard conditions (37 °C, 5% CO_2_). Culture medium was prepared as a 1:1 mixture of DMEM/F12 and Neurobasal medium (Gibco, #12348-017) supplemented with 10 ng/mL epidermal growth factor (EGF), 20 ng/mL fibroblast growth factors (FGF), 1X N-2 and 1X B-27 supplements, 1X Penicillin/Streptomycin, 1X Sodium-Pyruvate, 1X GlutaMAX, 1X MEM-NEAA, 4 mg/mL Heparin, 5 mM HEPES, 0.01 mM 2-Mercaptoethanol and 100 mg/L BSA. Differentiation of NSCs was induced with medium deficient of EGF and FGF, supplemented with 2% FBS (Sigma Aldrich, #F7524). NSCs were considered neurogenic in the first 5 DIV and gliogenic for the following passages as described before (Hirabayashi *et al*., 2009). Differentiation assays of neurogenic NSCs were performed at 2 DIV and of gliogenic NSCs after passaging every 3–4 day at 20 DIV. For immunohistochemistry analysis, NSCs were plated on poly-D-lysin- and laminin-coated coverslips (Marienfeld Superior, #0111520) in a 24-well plate, with either 10,000 cells per well for the NSC proliferation assay or 50,000 cells per well for the differentiation assay. Medium was changed every second day. Deletion of *Pcgf* genes was induced by addition of 1 μM 4-hydroxytamoxifen (OHT) to culture medium for 24–48 hours, as previously described (Fursova *et al*., 2019). 4-hydroxytamoxifen was dissolved in EtOH at a concentration of 1mM and stored for a maximum of 4 months at –20°C.

#### Immunohistochemistry analysis of tissue sections

Tissue was fixed in 4% PFA in 120 mM phosphate buffer pH 7.4 for 24 hours at 4°C, washed twice in PBS, transferred to 30% sucrose for 24 hours, embedded in O.C.T. compound (Sakura Finetek, #4583) with 15% sucrose and frozen on dry ice. Tissue was cut into 12 µm cryosections on a Thermo Fisher NX70 cryostat. Immunofluorescence was performed as previously described (Cubillos *et al*., 2024; Florio *et al*, 2015). Antigen retrieval for 1 hour with 10 mM citrate buffer pH 6.0 at 70°C in a water bath was followed by three washes with PBS, quenching for 30 min in 0.1 M glycine in PBS and blocking for 30 min in blocking solution (10% horse serum and 0.1% Triton in PBS) at room temperature. Primary antibodies were incubated in blocking solution over night at 4°C. Subsequently, sections were washed three times in PBS, incubated with secondary antibodies (1:1,000) and DAPI (1:1,000) in blocking solution for 1 hour at room temperature, and washed again three times in PBS before mounting on microscopy slides with Mowiol.

Images were acquired with a Zeiss ApoTome2 fluorescence microscope with a 20x objective using 1.5-µm thick optical sections. ZEN software was used to generate maximum intensity projections. To quantify the intensity of PRC1 markers, the human foetal and mouse embryonic tissue was divided into the different germinal zones and cortical plate based on SOX2 and CTIP2 staining and alignment of cells in DAPI within a 100 μm-wide image. The Fiji plug-in Stardist 2D (Schmidt *et al*, 2018) was applied using the versatile model to segment nuclei in DAPI, and if required, segmentation was corrected manually. Mean grey values of segmented nuclei were measured in Fiji and the resulting data processed using Excel and Prism software. Normal distribution of data was tested by Kolmogorov-Smirnov and Shapiro-Wilk tests, followed by Tukey’s multiple comparison test.

#### Immunohistochemistry analysis of NSC differentiation

Cells were fixed in 2% PFA in 120 mM phosphate buffer pH 7.4 for 10 min at room temperature, before PFA was washed away twice with PBS. Cells were permeabilized with 0.1% Triton in PBS for 5 min, followed by washing twice with PBS for 5 min and twice with washing solution containing 0.1% Tween in PBS for 5 min. After this, cells were blocked for 30 min in blocking buffer containing 2.5% BSA (Sigma Aldrich, #A2153), 0.1% Tween and 10% horse serum in PBS, before primary antibodies diluted in blocking buffer were added and incubated over night at 4°C. This was followed by three washes with washing solution for 10 min, incubation with secondary antibodies (1:1,000) and DAPI (1:1,000) in blocking buffer for 1 hour at room temperature, three additional washes and embedding in a drop of Mowiol on microscopy slides.

Images were acquired with a Zeiss ApoTome2 fluorescence microscope with a 20x objective using 1.5-µm thick optical sections. ZEN software was used to generate maximum intensity projections. Samples were blinded after staining, before acquisition of images. For quantification of marker positive cells, nuclei were segmented in DAPI using the Fiji plug-in Stardist 2D (Schmidt *et al*., 2018) and counted with the cell counter tool in Fiji. The resulting data was processed using Excel and Prism software. Data was analysed for outliers using the Grubb’s test, followed by Tukey’s multiple comparison test.

#### Protein expression analysis by Western blotting

Proteins were isolated from cells in culture using TOPEX Plus buffer containing 300 mM NaCl, 50 mM Tris-HCl pH7.5, 0.5% Triton, 1% SDS, 1 mM DTT (Roche, #10708984001), 1X protease inhibitor (Roche, #4693116001) and 333.33 U/mL Benzonase (Sigma Aldrich, #E1014-25KU) in water, by incubation at room temperature until viscosity disappeared (5–15 min). Protein concentration was measured using the Pierce detergent compatible Bradford assay kit (Thermo Scientific, #23246). Subsequently, 40 µg of protein were resolved on a 10% SDS Polyacrylamide gel and transferred to a PDVF transfer membrane (ThermoScientific, #88518). Membranes were blocked for 1 hour at room temperature with 5% skim milk in PBS with 0.05% Tween, and then incubated with primary antibodies over night at 4°C. Secondary antibodies were incubated for 1 hour at room temperature. Antibody signal was detected using the SuperSignal West Pico plus kit (ThermoFisher Scientific, #34579).

#### RNA-seq library preparation

RNA-seq was performed as previously described (Cubillos *et al*., 2024). RNA of differentiating NSCs was isolated using the Quick-RNA MiniPrep kit (Zymo Research, #R1008). Transcriptome libraries were prepared using an adapted version of the SmartSeq2 protocol (Picelli *et al*, 2013). Isolated total RNA from an equivalent of one 24-well was denatured for 3 minutes at 72°C in 4 μL hypotonic buffer (0.2% Triton-X 100) in the presence of 2.4 mM dNTP, 240 nM dT-primer and 4 U RNase Inhibitor (NEB, M0314L). Reverse transcription was performed at 42°C for 90 min after filling up to 10 µL with RT buffer mix for a final concentration of 1X Superscript II buffer (Invitrogen), 1 M Betaine, 5 mM DTT, 6 mM MgCl2, 1 µM TSO-primer, 9 U RNase inhibitor and 90 U Superscript II. The reverse transcriptase was inactivated at 70°C for 15 min. For subsequent PCR amplification of the cDNA, the optimal PCR cycle number was determined with an aliquot of 1 μL unpurified cDNA in a 10 μL qPCR containing 1X Kapa HiFi Hotstart Readymix (Roche), 1X SybrGreen and 0.2 μM UP primer. The residual 9 μL cDNA were subsequently amplified using Kapa HiFi HotStart Readymix (Roche) at a 1X concentration together with 250 nM UP-primer under the following cycling conditions: initial denaturation at 98°C for 3 min, 22 cycles [98°C 20 sec, 67°C 15 sec, 72°C 6 min] and final elongation at 72°C for 5 min. Amplified cDNA was purified using 1X volume of Sera-Mag SpeedBeads (GE Healthcare) resuspended in a buffer consisting of 10 mM Tris, 20 mM EDTA, 18.5% (w/v) PEG 8000 and 2 M sodium chloride solution. The cDNA quality and concentration were determined using a Fragment Analyzer (Agilent).

For library preparation, 2 µL amplified cDNA was tagmented in 1X Tagmentation Buffer using 0.8 µL bead-linked transposome (Illumina DNA Prep, (M) Tagmentation, Illumina) at 55°C for 15 min in a total volume of 4 µL. The reaction was stopped by adding 1 µL of 0.1% SDS (37°C, 15 min). Magnetic beads were bound to a magnet, the supernatant was removed, beads were resuspended in 14 µL indexing PCR Mix containing 1X KAPA Hifi HotStart Ready Mix (Roche) and 700 nM unique dual indexing primers (i5 and i7), and subjected to a PCR (72°C 3 min, 98°C 30 sec, 12 cycles [98°C 10 sec, 63°C 20 sec, 72°C 1 min], 72°C 5 min). Libraries were purified with 0.9X volume Sera-Mag SpeedBeads, followed by a double size selection with 0.6X and 0.9X volume of beads, and a final 0.9X purification to obtain a fragment size distribution of 200–700 bp. Sequencing was performed after quantification using a Fragment Analyzer on an Illumina Novaseq 6000 in 100 bp paired-end XP mode with an average sequencing depth of 40 million fragments per library.

#### RNA-seq data analysis

Quality control of the sequencing data was performed with FastQC (version 0.11.9). Kallisto (version 0.64.1) (Bray *et al*, 2016) was used to align trimmed reads to mouse GRCm39. For further processing, data was imported into R using Tximport (Soneson *et al*, 2015) and EnsDb.Mmusculus.v79 packages. Raw fragment normalization based on library size and testing for differential expression between genotypes was performed with DESeq2 (version 1.30.1; Wald test) (Love *et al*, 2014) with a false discovery rate (FDR) of 5% and a Log2 fold change threshold of 1. To quantify gene expression levels within samples, transcripts per million (TPM) values were calculated with Tximport (Soneson *et al*., 2015). Volcano plots and heatmaps were generated with the R-packages Ggplot2 (Wickham *et al*, 2016) and Complexheatmap (Gu *et al*, 2016). Venn Diagrams were generated using the online platform DeepVenn.com (Hulsen, 2022). Differentially expressed genes were analysed by KEGG pathway (https://www.genome.jp/) and gene ontology (https://geneontology.org/) (Ashburner *et al*, 2000; Thomas *et al*, 2022) analysis.

#### Statistical analysis

Sample sizes are reported in each figure legend. Sample sizes were estimated based on previous literature (Albert *et al*., 2017; Cubillos *et al*., 2024; Schmitz *et al*., 2011). All statistical analysis was performed using Prism (GraphPad Software). Normal distribution of datasets was tested by Shapiro-Wilk or Kolmogorov-Smirnov test. Data was analysed for outliers using the Grubb’s test. The tests used included Student’s *t*-test and Tukey’s multiple comparison test, as indicated in the figure legend for each quantification. Significant changes are indicated by stars for each graph and described in the figure legends.

## Acknowledgements

We are grateful to the facilities of the CRTD and DRESDEN-concept partner institutions for the outstanding support provided, notably R. Hans from the Light Microscopy Facility, the teams for animal husbandry and histology and the DRESDEN-concept Genome Center lab team for technical support. We thank all members of the Albert laboratory for help and discussions. We acknowledge R. Kageyama from Kyoto University for providing the *Nes*::CreERT2 mouse line. MA acknowledges funding from the Center for Regenerative Therapies TU Dresden, the DFG (Emmy Noether, AL 2231/1-1) and the Schram foundation.

## Author contribution

**Janine Hoffmann:** Conceptualization; investigation; formal analysis; visualization; writing – review & editing; **Theresa M. Schütze:** Formal analysis, writing – review & editing; **Annika Kolodziejczyk:** Investigation; resources; writing – review & editing; **Annekathrin Kränkel**: Investigation; **Susanne Reinhardt**: Resources; writing – review & editing; **Razvan P. Derihaci:** Resources; **Cahit Birdir:** Resources; **Pauline Wimberger:** Resources; **Haruhiko Koseki**: Resources; **Mareike Albert:** Conceptualization; writing – original draft; writing – review & editing; funding acquisition; supervision.

